# Astral architecture can enhance mechanical strength of cytoskeletal networks by modulating percolation thresholds

**DOI:** 10.1101/2025.06.17.660175

**Authors:** Brady Berg, Jun Allard

## Abstract

A repeated pattern in cytoskeletal architecture is the aster, in which a number of F-actin filaments emerge star-shaped from a central node. Aster-based structures occur in cytoplasmic actin, the early stages of the cytokinetic ring in yeast, and in the context of biomimetic materials engineering. In this work, we use computational simulation to show that there is an optimal number of filaments per aster that maximizes rigidity, even at a fixed density of F-actin. This nonlinear dependence holds for both the shear and extensional moduli. At physiological parameters, the maximum corresponds approximately to the same filaments-per-aster observed in recent super-resolution images of cortical F-actin. Furthermore, we find that increasing filaments-per-aster leads to dramatic increases in the sample-to-sample variability in network rigidity. We explain both effects using percolation theory, wherein the probability that a given network is productively connected exhibits a sharp dependence on parameters. The dependence of network rigidity on this nanoscale architectural feature may suggest a mechanism by which cells tune the physical properties of their actin networks locally and rapidly (since no new F-actin must be assembled) and may inform efforts to create adaptive synthetic metamaterials inspired by actin networks.

## Introduction

As the key component of cell mechanical structures (including many approximately 2-dimensionsal structures, e.g., the cortex [1–3] and the cytokinetic ring [4]), the mechanical properties of F-actin are crucial to understanding cell biology [5, 6]. Much progress has been achieved understanding F-actin at the scale of individual filaments [7, 8], and at the scale of cells, where F-actin forms a cytoskeleton that can be understood as an active gel using continuum models [9]. Connecting those two scales is the nanoscale architecture — that is, the precise arrangement of 10s-1000s of individual filaments, and how these give rise to the bulk properties that provide the starting point for active gel theory. Nanoscale architecture has been studied in physical theory and computational simulations [10–17]. With the advent of super-resolution microscopy, an increasing number of actin structures have been experimentally studied, for example the low-density cytosolic network [18, 19] and the cortex [20]. Several recent works have demonstrated that classical percolation theory is useful in understanding the actin cytokeleton and other filamentous structures [10, 21–26].

A major question is how cells rapidly tune the nanoscale architecture of their F-actin cytoskeleton to achieve responsive cell-wide mechanical properties. Recently, García-Arcos et al. [26] observed that cells locally and rapidly alter the mechanical properties of the cytoskeleton during motility. The reserchers found that cells explore a phase diagram where F-actin density and molecular motors give rise to mechanical properties, with non-monotonic phase transition lines [26].

A recurring observation in vivo [18–20], in vitro [27], and in computational simulations [27, 28] is the emergence of asters, in which several actin filaments emanate from an astral center in a star-like pattern as shown in Figure 1. Kruse et al. [29] provide evidence that asters are a ubiquitous, fundamental, elementary unit of cellular organization. Actin asters are varied in structure, with centers containing formin [18] or Arp2/3 [20], either requiring myosin [27] or not [20]. In vivo, the precise quantification of asters remains challenging, but recently, Xia et al. [20] used super-resolution microscopy to find that the cortex of embronic stem cells was composed of asters with *a*_*n*_ ≈ 3 − 6 filaments per aster, what they term the “astral coordination number” and we herein call “astral number” *a*_*n*_. Much prior work has queried how asters may emerge [27, 28]. Here instead we ask, what are the biophysical consequences of asters? Note that these random glassy configurations with rigid crosslinks give rise to more complexity than classical Maxwell constraint counting argument [30] would suggest (see, e.g., [23, 31]).

**Figure 1:**
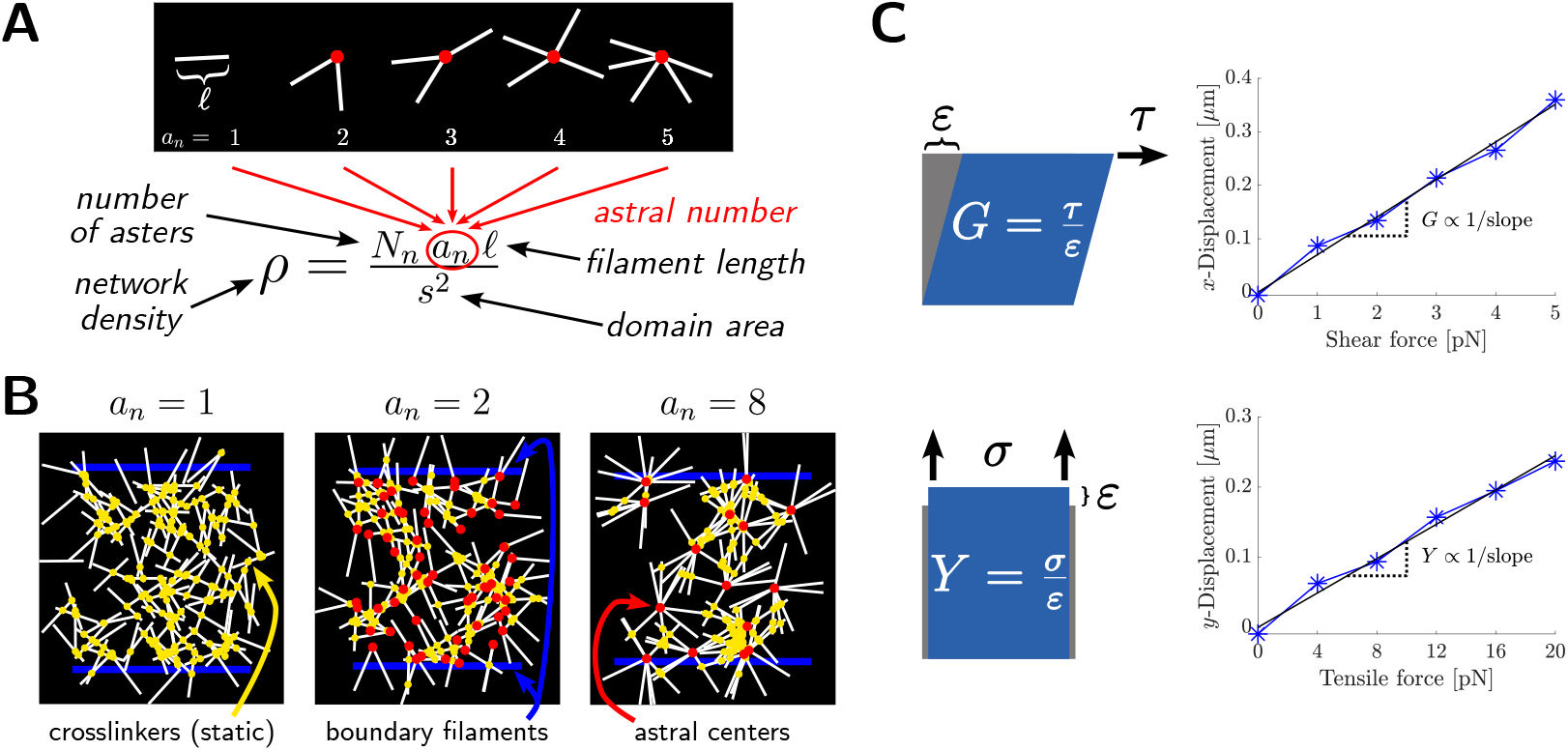
Framework for studying astral network rigidity. (a) Schematic of asters and their astral number *a*_*n*_, i.e. filaments per aster. Variations in *a*_*n*_ at a fixed network density *ρ* correspond to changes in the nanoscale arrangement of a set number of filaments. (b) Schematics of filament networks at fixed density *ρ* and astral numbers 1 (left), 2 (middle), and 8 (right). Crosslinkers are shown in yellow, boundary filaments are shown in blue, and astral centers are shown in red. (c) Schematics of bulk shear modulus *G* (top) and bulk Young’s modulus *Y* (bottom). We estimate these elastic moduli from the slope of force (proportional to stress) vs. displacement (proportional to strain) simulation data.

In this work, we simulate a minimal computational model of astral polymer networks, and measure their mechanical strength. The model takes as input the formation of asters with a particular astral number, linked with additional inter-aster crosslinks. The simplicity of this model allows dissection of the consequences of specific architectural features, similar to the way in which in vitro reconstitution systems have aided our understanding of the actin cytoskeleton [25, 32, 33]. We find an optimal astral number that maximizes the mechanical moduli, for a fixed total amount of polymer. Furthermore, the astral number that maximizes mechanical strength is in close agreement with the aster coordination number of Xia et al. [20], suggesting that the cells under their study were optimizing nanoscale architecture to maximize strength for a given amount of F-actin. Our results — and the nonlinear phase diagram of García-Arcos et al. [26] — emerge from critical percolation thresholds.

## Results

### Computational model of astral filament network

To understand the properties of astral cytoskeletal networks, we extended the model of Wilhelm and Frey [10] to the case of astral network components. Asters are represented in the model as radial assemblies each with *a*_*n*_ filaments per central node (their *astral number a*_*n*_, see Figure 1a), each filament having fixed length ℓ. Every aster in a particular network has the same number of filaments per central node. To form a network, *N*_*n*_ asters were distributed in a square domain of area *s*^2^ and permanently binding crosslinkers were distributed to crosslink asters to each other and to the boundary at the top and bottom of the domain (Figure 1b). Coordinates of astral centers and the orientation of each filament about its respective center were sampled uniformly at random. We use network density *ρ* to refer to the average filament length per unit network area,

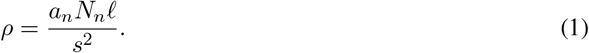

Unless otherwise specified, we use ℓ = 1 µm and *s* = 10 µm; for a complete list of default parameter values see Table 1.

**Table 1:** Biophysical parameters.

| Parameter (symbol) | Default value | Description |
| --- | --- | --- |
| system size ($s$) | $10\text{ }\mu\text{m}$ | side length of square domain |
| network density ($\rho$) | $7.5\text{ }\mu\text{m}^{-1}$ | average filament length per unit area |
| filament length ($\ell$) | $1\text{ }\mu\text{m}$ | inextensible; no growth/disassembly/fracture |
| filament bending rigidity ($k_{\text{bend}}$) | $20\text{ pN }\mu\text{m}^2$ | |
| astral center stiffness #1 | $1000\text{ pN }\mu\text{m}^{-1}$ | attaches filament ends to center of aster |
| astral center stiffness #2 ($k_{\text{rot}}$) | $500\text{ pN }\mu\text{m}^{-1}$ | resists filament rotation about center of aster; acts $0.3\text{ }\mu\text{m}$ from center |
| boundary filament bending rigidity | $1 \times 10^3\text{ pN }\mu\text{m}^2$ (shear);<br>$1 \times 10^6\text{ pN }\mu\text{m}^2$ (tensile) | |
| boundary anchor stiffness | $1 \times 10^3\text{ pN }\mu\text{m}^{-1}$ | fix lower boundary filament in place (and upper boundary during crosslinking phase) |
| boundary anchor spacing | $1\text{ }\mu\text{m}$ | anchors placed along lower boundary filament |
| crosslinker concentration | $\approx 225\text{ particles }\mu\text{m}^{-2}$ | see Materials and Methods text for details |
| crosslinker stiffness | $100\text{ pN }\mu\text{m}^{-1}$ | |
| crosslinker binding rate | $10\text{ s}^{-1}$ | |
| crosslinker unbinding rate | 0 | crosslinkers do not unbind |
| crosslinker diffusivity | $10\text{ }\mu\text{m}^2\text{ s}^{-1}$ | |
| crosslinker binding range | $0.01\text{ }\mu\text{m}$ | |
| shear force range | 0 pN to 5 pN | number of equally spaced force values simulated per network specified in each figure caption |
| tensile force range | 0 pN to 20 pN | number of equally spaced force values simulated per network specified in each figure caption |
| viscosity | $0.01\text{ pN s}/\mu\text{m}^2$ | |

### Rigidity of astral filamentous networks exhibits a maximum at intermediate astral number

We next simulated networks for varying astral number *a*_*n*_ and density *ρ* (Figure S2). We quantified the rigidity of these astral networks by estimating an elastic modulus from the slope of stress-strain response curves.

To do this, we applied a varying force to each sampled network, computed center-of-mass displacements at steady state, and estimated elastic modulus from the initial linear response (Figure 1c). Modulus estimation is detailed further in Materials and Methods. We investigated network behavior under shear and under tension. Representative images of networks experiencing a varying shear force are provided in Figure 2a.

**Figure 2:**
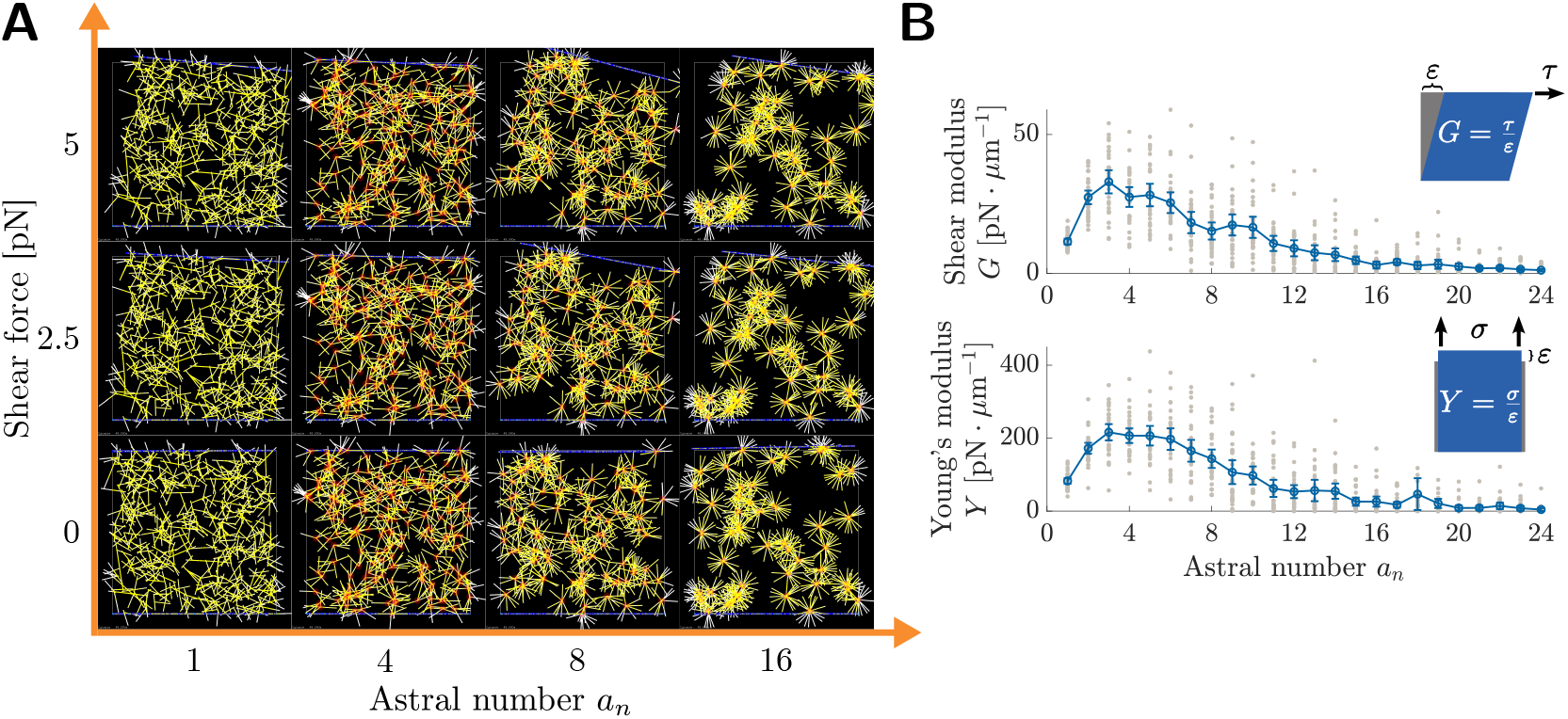
Rigidity of astral filamentous networks exhibits a maximum at intermediate astral number, even at fixed density. (a) Steady-state snapshots of astral network deformation in response to applied shear forces. All tiles have equal total number of filaments. (b) Elastic moduli of astral networks at density *ρ* = 7.5 µm^−1^ as a function of astral number. Upper: shear modulus, lower: Young’s (tensile) modulus. Plots show mean and 95% CIs from *N* = 30 network samples using 6 force values per network (see Materials and Methods for details). Individual network moduli are shown (gray markers).

We observed a peak in both the shear and Young’s (tensile) moduli at an intermediate astral number (Figure 2b). These peaks in modulus occur at similar astral numbers for both deformation modes. The location of such peaks is insensitive to the specific network density tested (Figure S2). The peak location also does not depend on crosslinker density or crosslinker stiffness, and the modulus values themselves appear to scale similarly with both quantities (Figure S3). The peak is not sensitive to the particular filament bending rigidity selected, though at especially low bending rigidities (~1 pN µm^2^) networks trend toward lower modulus values (Figure S4a). The presence of angular stiffness (i.e. torques) at astral centers is not necessary to observe a peak in modulus values (Figure S4b).

### Weakening at high astral number is due to increasing probability of network failure

The sample-to-sample variability in elastic modulus (bars in Figure 2b) also depends on aster number. Whereas for non-astral networks (*a*_*n*_ = 1) the variance in moduli is low, at moderate astral number the spread of the sampled moduli values appears much larger. This suggests that network rigidity is more sensitive to the initial configuration of network components when astral number is higher. To investigate further the role of geometric variability in determining network rigidity, we estimated the distributions of the shear and Young’s moduli for various astral numbers (Figure 3a). As in Figure 2b, the distributions for shear and Young’s moduli have similar shapes, but this shape changes categorically depending on aster number. For non-astral networks (*a*_*n*_ = 1), the distributions of moduli have vanishing probability of very small moduli, i.e., their histograms sit away from the origin and the cumulative distribution functions (CDFs) are flat at the origin (purple CDFs and histograms in Figure 3a), like a Gaussian distribution. Once astral number increases past *a*_*n*_ ≈ 4, the distributions become increasingly skewed towards low modulus values, losing the upward concavity at zero by *a*_*n*_ = 8 and switch to positive probability of small moduli by *a*_*n*_ = 16 (gold CDFs and histograms in Figure 3a), like an exponential distribution.

**Figure 3:**
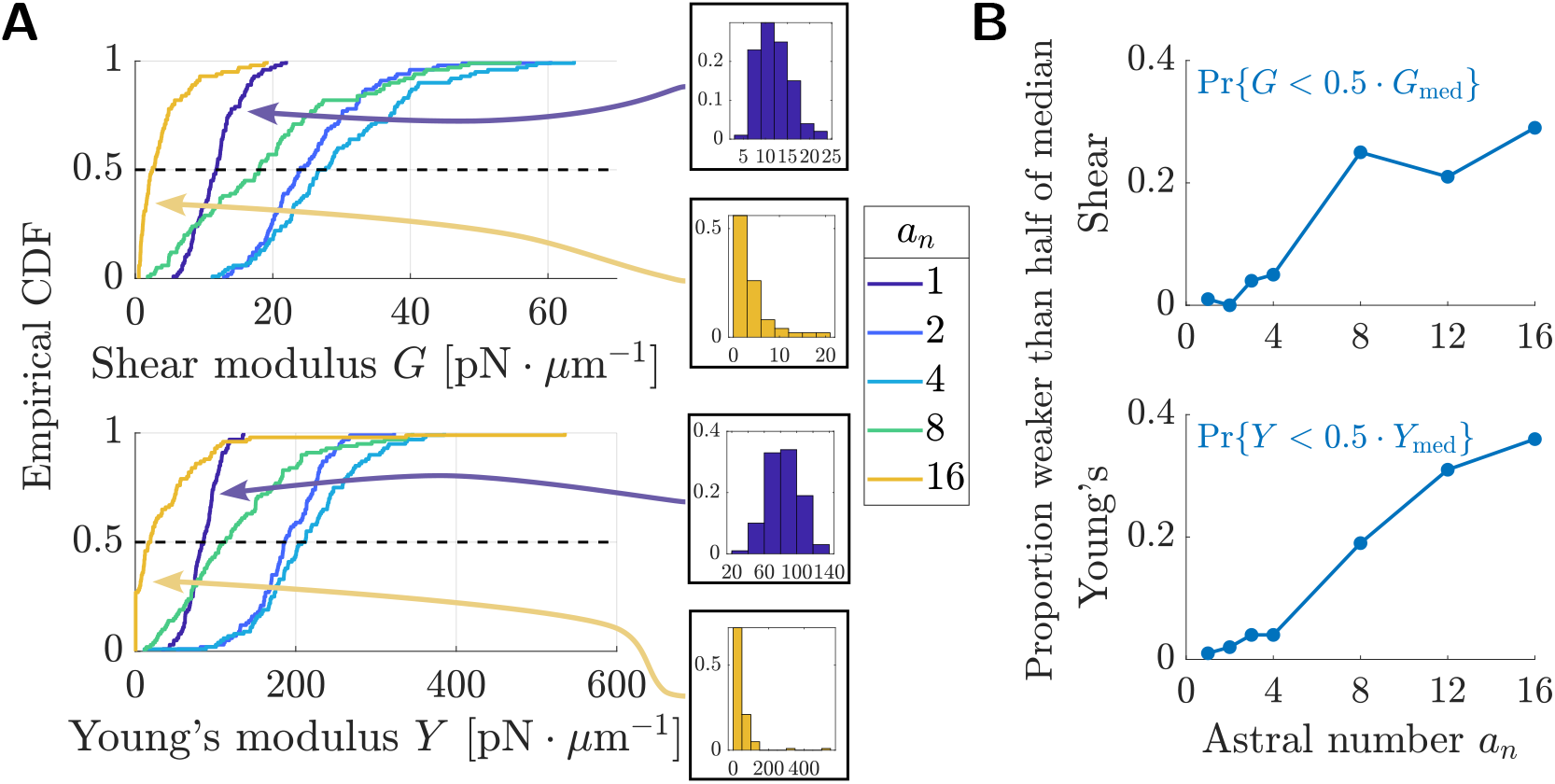
Weakening at high astral number is due to increasing probability of network failure. (a) Cumulative distribution functions and histograms for the elastic moduli of astral networks at *ρ* = 7.5 µm^−1^. Upper: shear modulus, lower: Young’s (tensile) modulus. (b) Proportion of astral network that are weaker than half the median modulus. Distributions were estimated from *N* = 100 network samples using 4 force values per network (see Materials and Methods for details).

This marked shift from Gaussian-like to exponential-like distribution types suggests that increasing *a*_*n*_ beyond intermediate values results in a bias towards weak and mechanically incompetent networks at these astral numbers. We computed, at each sampled astral number, the proportion of networks which had modulus below half of the respective median modulus value (Figure 3b). For astral numbers below ≈ 4, the proportion of networks which are weak relative to the median is less than 5%. This quantity dramatically increases, approaching 30 − 40% at high astral numbers, even as the median modulus at high astral numbers itself decreases (intersections with dashed lines in Figure 3a). Thus the weakening at high astral numbers is driven in part by an increase in the likelihood of a network geometry being incapable of withstanding mechanical load (e.g., due to fractures or gaps).

### Strength dependence on astral number is not associated with dangling filament fraction nor mean segments per node

One *a priori* possible explanation for why networks with intermediate astral number are more rigid than non-astral networks is that the astral centers force more of the available filament length into mechanically productive structures. One way to classify if a filament segment is mechanically productive is to distinguish between filament segments which are bounded on both ends by a connection (either astral center, or inter-aster crosslink, shown as orange segments in Figure 4a) and those which are not; those in the latter category are thus dangling ends (black segments in Figure 4a). Dangling ends do not experience any net force when a network is deformed.

**Figure 4:**
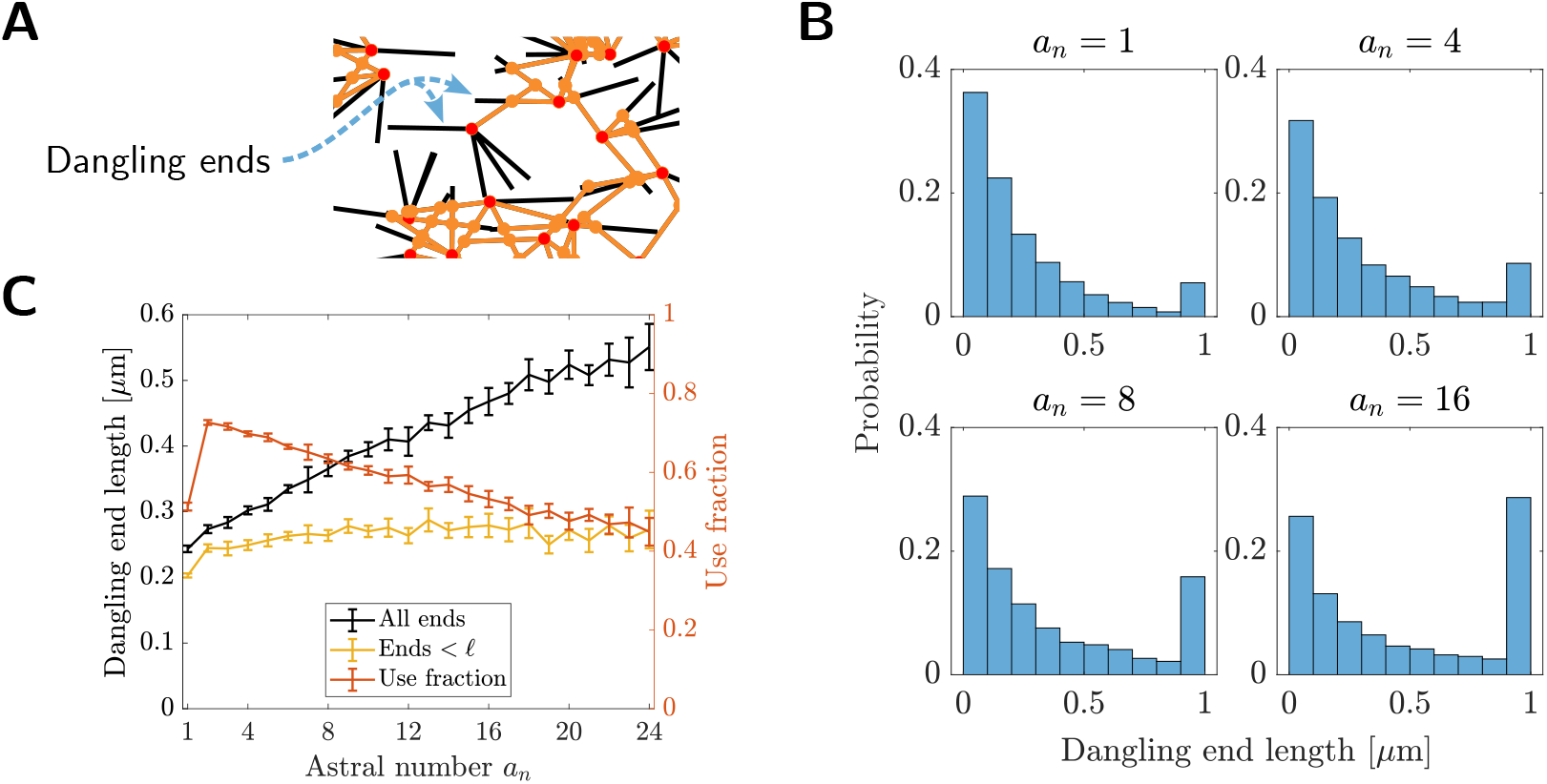
Strengthening over low astral numbers is not associated with dangling filament fraction. (a) Schematic of dangling ends (black sections of filaments) in astral networks. (b) Histograms of dangling end lengths at various astral numbers. Distributions are aggregated from the dangling ends in 10 networks at each astral number. (c) Mean dangling end length as a function of astral number. Also shown is the length “use-fraction”, defined in Equation 2. Bars indicate 95% CIs estimated from 10 networks.

To test if changes in dangling ends could explain the rigidity increase shown in Figure 2b, we measured the dangling end lengths in networks with different astral numbers (see Materials and Methods for details). Distributions of dangling ends have the shape of an exponential distribution with a singular accumulation point corresponding to the maximum dangling end length ℓ = 1 µm (Figure 4b). At sufficiently high astral numbers, this accumulation point gathers enough probability that the distributions appear bimodal. The exponential part of these distributions is in agreement with analytic studies which predict the distribution of dangling ends to be exponential [34].

The mean dangling end length in astral networks increases monotonically with *a*_*n*_ (Figure 4c). This increase vanishes when inspecting only the dangling ends with lengths strictly below ℓ, showing that the change in mean dangling end length is driven by the number of totally isolated filaments. As a complimentary nquantification, we computed the length “use-fraction” for all sampled networks, defined as

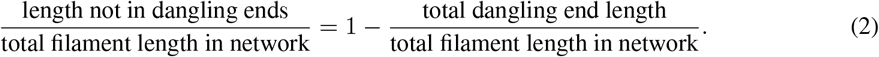

While the use-fraction increases when *a*_*n*_ changes from 1 to 2, it decreases consistently for *a*_*n*_ *≥* 2. The initial increase in use fraction is because there are fewer total dangling ends in astral networks, since filament ends located at astral centers are considered to be crosslinked to each other precisely at their ends. (In non-astral networks, both ends of a filament generate dangling ends of nonzero length.) Overall, the trends pertaining to dangling ends in astral networks does not agree with the trends in modulus in Figure 2b.

Classical rigidity analysis [30] suggests that the mean number of segments per node plays a role in determining deformability. While the networks studied here are more complex [23, 31], we compute the mean segments per node, and its full distribution, as a function of aster number, in Figure S5. There are two kinds of nodes between filaments in this model: astral centers, which all have fixed *a*_*n*_ filaments each, and then inter-aster crosslink connections. The inter-aster crosslinks connect either 2, 3 or 4 productive segments, where two line segments may belong to the same filament. We find that the mean number of segments (for both kinds of nodes) gradually increases from around 3.1 to around 3.9 as astral number increases from *a*_*n*_ = 1 to 24, without an optimum or crossing integer values.

### Mechanical rigidity is associated with network connectedness

Having confirmed that dangling ends and use-fraction do not explain the maximum in astral number, we next noted that at high astral numbers, filaments are highly concentrated around relatively few nodes, making it extremely unlikely that network components connect into a structure capable of resisting force. Following this, we sought to study the connectivity properites of astral networks by quantifying their percolation thresholds, a geometric measure which has previously proven useful in understanding the actin cytokeleton and other filamentous structures [10, 21–26]. In brief, the central idea of percolation theory is that as network density increases, the probability that a random network is connected transitions sharply from 0 to 1. The density at which this transition occurs is known as the percolation threshold. Though we restrict our attention to finite networks in this work, the behavior of finite systems is similar and approaches the infinite size for *s ≫* ℓ (Figure S6).

We computed two different percolation probabilites for finite-size astral networks (ℓ = 1 µm, *s* = 10 µm) via Monte Carlo sampling (see Materials and Methods for details). Given the mode of external stress we are interested in, we define two distinct notions of connectedness and therefore percolation. First, we investigated the appearance of *spanning* components in astral networks, defined as a set of connected asters that contains a crosslink both above and below the two system boundaries (Figure 5a). The existence of a spanning component is sufficient for an astral network to withstand applied force. Second, we investigated the emergence of a *unique* connected component (Figure 5b), i.e. when all the asters in a network crosslink into a single bulk material. We refer to this percolation mode as *connectivity* percolation. The computed percolation probabilites are shown in Figure 5c. As expected, for either definition, the probability transitions rapidly, and there is only a small region of parameters with subpopulations of both connected and unconnected networks.

**Figure 5:**
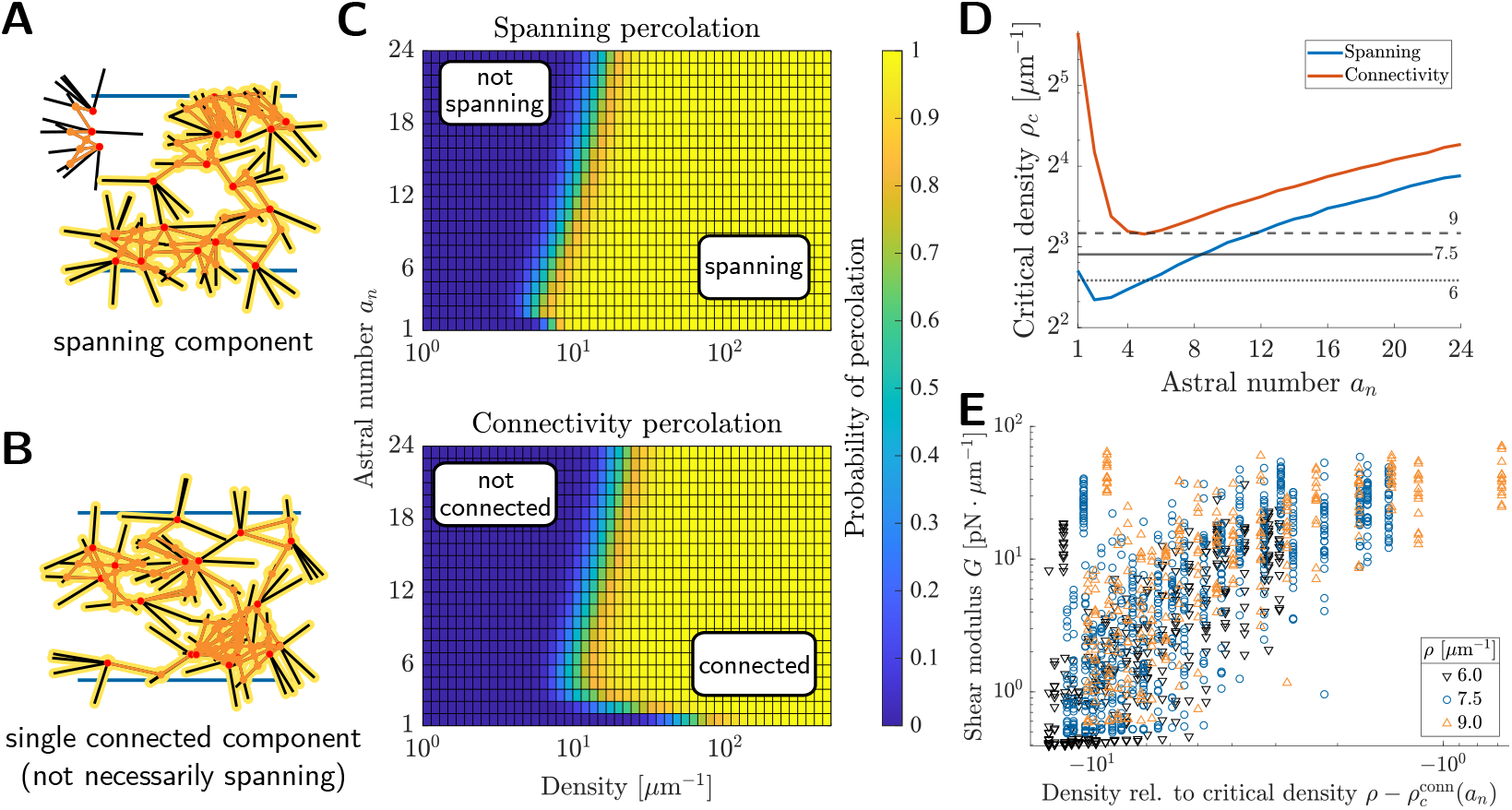
Mechanical rigidity is associated with network connectedness. (a) Schematic of connected component that spans between the top and bottom edges of the domain. (b) Schematic of a network where all asters are connected into a single (unique) component. (c) Heatmaps showing percolation probabilities for astral networks with ℓ = 1 µm and *s* = 10 µm. Probabilites were estimated from *N* = 2000 networks at each sampled density and astral number. (d) Critical percolation densities estimated from the data in (c) as a function of astral number. The critical percolation density is defined as the filament density where the percolation probability reaches 0.5. (e) Shear modulus versus 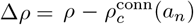, the difference between the filament density in the simulated network *ρ* and the critical connectivity percolation density 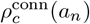. Three network densities are shown (see also Figure S2).

Notably, the percolation thresholds appear to be non-monotonic in astral number. We estimated these thresholds via computing the density where the percolation probability reaches 0.5, henceforth referred to as the critical densities 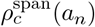 or 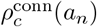. (Note that these quantities are approximations for the infinite-extent percolation threshold as *s → ∞* as shown in Figure S6.) These critical densities are plotted in

Figure 5d. While both curves exhibit a minimum at low but nonzero astral number, we observed that the region of optimality in the connectivity threshold curve corresponds to the region of optimality in Figure 2b. We thus speculated that the proximity of the network density to the critical connectivity density was a predictor of network rigidity. To test this, we aggregated shear modulus samples from three different network densities *ρ* and various *a*_*n*_ (see also Figure S2) and plotted shear modulus against the distance to criticality 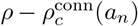 (Figure 5e). We see a clear correlation between rigidity and network density relative to 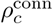 across astral number and network density (Pearson correlation of log-scale data 0.5419). This suggests that the dependence of rigidity on astral number may be due in part to the shifting of the critical connectivity densities.

## Discussion

In a seminal work, Wilhelm and Frey [10] studied a simple model with only two molecular components: semi-flexible filaments and inter-filament crosslinkers, in 2 dimensions. They introduced the name “Mikado model” in reference to a children’s game with similar architecture, and found nontrivial behavior including a new universality class of mechanical response. Continuing this agenda, we study a simple model with three molecular components: inextensible filaments, mechanical linkers creating astral centers, and mechanical linkers creating inter-aster crosslinks, thus creating an “astral Mikado model”. In both cases, the simplicity allows the dissection of the specific role of model features, for example by disentangling the cause of astral architectures with the consequence of astral architecture, and by disentangling bulk (passive) stress response from active contractility. The simplicity also allows efficient simulation of large systems over many parameters, which would be computationally expensive in models with more features [35, 36].

The major finding of this work is an optimal astral number that maximizes mechanical strength, both shear modulus and Young’s modulus, for a fixed density of F-actin filaments. As other parameters are varied, like density of F-actin, the parameter dependence of mechanical moduli are nonlinear and non-monotonic, but we find the dependency largely aligns with a single metric, the density difference with the percolation threshold density (Figure 5e). This suggests that the strength of the network is a consequence of the heterogeneity in local filament density, which creates gaps in the network, rather than the constraints of different numbers of filaments and joints as in classical Maxwell counting [30].

In terms of problem-solving strategies selected by cells, a maximum rigidity may be desirable in some cases for cell integrity [37], and undesirable in other cases, for example during cell migration through a fine mesh [38, 39]. So, how does the astral number of in vivo F-actin networks compare with the optimal number we found here? Xia et al. [20] used super-resolution microscopy to study the F-actin cortex in embryonic stem cells. They identifies astral centers (excluding inter-astral crossovers by filtering by the molecular constituents of astral centers), and then counted the filaments emanating from these. They found *a*_*n*_ ≈ 4 with an inter-quartile range of 3 − 5, in remarkable agreement with the optimum identified by the astral Mikado model (Figure 2). This astral number is also roughly consistent with images of Luo et al. [18] and quantification in Vavylonis et al. [4], in different cellular contexts. Thus, this may provide an example where a single biophysical parameter optimizes a clearly-defined objective function in a cell biological system. Indeed, synthetic aster-based materials have been studied as a possible meta-material [40–42].

Recently, García-Arcos et al. [26] demonstrated that migrating cells locally and dynamically tune their mechanical properties to navigate complex environments, and inferred a phase diagram depending on F-actin density and myosin contractility (their Figure 5E). While we do not explicitly consider myosin contractility, Wollrab et al. [27] demonstrated both in vivo and in silico that myosin organizes F-actin into astral structures, in agreement with other work [28]. Thus, if we assume that myosin activity causally induces astral structure, then the vertical axis of the phase diagram we show in Figure 5c roughly corresponds to myosin contractility. With this correspondence, the phase diagram in Figure 5c is remarkably similar to the phase diagram inferred by [26]. This suggests that dynamic astral nanoscale architecture may provide the knob allowing cells to dynamically tune their mechanics.

Future models of astral cytoskeletal networks will explore polydispersity in astral number and in filament length. Since many distributions can correspond to the same mean astral number, it would be interesting to examine how properties of the distribution (e.g. number of aster types, variance) modify the trends seen here. Regarding filament length, previous investigations [34] demonstrated a role for filament length heterogeneity. The rigidity of such networks appears to be sensitive to the presence of long filaments both experimentally [20] and computationally [34], and modeling can explore how astral nanostructure magnifies (or diminishes) this sensitivity.

## Materials and Methods

### Mechanical simulations

Mechanical simulations of astral cytoskeletal networks were carried out using a custom extension of Cytosim [43]^1^. Filaments were modeled as inextensible filaments of length ℓ = 1 µm and bending rigidity 20 pN µm^2^. Asters group filaments into radial assemblies via the action of two springs at the astral center with stiffnesses 1000 pN µm^−1^ and 500 pN µm^−1^. The former pins one end of a filament to the astral center, and the latter resists a filament’s rotation about the astral center. Orientations of filaments about each astral center were initialized uniformly at random.

Each astral network consists of a number of asters distributed uniformly at random within a square domain of side length *s* = 10 µm. All asters in a particular network have the same number of filaments per central node, and we use the term *astral number a*_*n*_ to refer to this quantity (see also Figure 1a). In addition to asters, two additional boundary filaments were placed at the top and bottom of the domain. Each boundary filament has length equal to *s*, the domain size, and rigidities either 10^3^ pN µm^2^ (shear force simulations) or 10^6^ pN µm^2^ (tensile force simulations).

Irreversibly-binding crosslinkers were distributed randomly over the domain area and have a binding radius of 0.01 µm, diffusivity of 10 µm^2^ s^−1^, and (linear) stiffness 100 pN µm^−1^. Crosslinkers do not individually exert torques. Unless otherwise indicated, the number of crosslinkers was 30× number of filaments, which at density 7.5 µm^−1^ corresponds to 225 particles µm^−2^. A high density of crosslinkers was selected to encourage binding between filaments before they have time to diffuse away from their initial positions, thus mimicking the simulations of Wilhelm and Frey [10] where crosslinks were placed wherever filaments intersect. Additional crosslinkers were placed near boundary filaments (10 particles µm^−1^ *· s* at each boundary) to ensure the connection of the astral network to these boundary filaments. Network components are schematized in Figure 1b. See Table 1 for a summary of the key biophysical parameters used in Cytosim; unless otherwise stated, parameters assume the indicated default value. Parameters related to the numerical scheme are summarized in Table 2.

**Table 2:** Cytosim simulation parameters.

| Parameter | Default value | Description |
| --- | --- | --- |
| time step | 0.01 s |  |
| numerical tolerance | 0.05 | default value, unitless |
| filament segmentation | 0.2 $\mu\text{m}$ | length of rigid filament sub-segments |
| boundary filament segmentation | 0.5 $\mu\text{m}$ | as above but for boundary filaments |
| duration of crosslinking phase | 5 s | system evolves without any applied force; initial network position recorded at end of this phase |
| duration of force application phase | 40 s | constant applied force; final network position recorded at end of this phase |

Each simulation begins by letting network components (asters and crosslinkers) diffuse freely for 5 s, allowing crosslinkers to bind asters together into a particular network geometry. During this time, the top and bottom boundary filaments are held in place. Since crosslinkers diffuse quickly relative to filaments and do not unbind, the network geometry that forms is dictated primarily by the random initialization of aster positions.

To study the response of each network to an applied force, we customized Cytosim to enable the application of force to the ends of particular network fibers. At *t* = 5 s, the position of each fiber in the network was recorded as the initial state of the network. Simultaneously, the top boundary filament was subjected to a constant applied force and the anchors holding it in place were removed. For shear force simulations, a rightward force was applied to the right-hand end of the top boundary filament. For tensile force simulations, upward forces were applied to each of the left- and right-hand ends of the top boundary filament, each with magnitude half of the reported applied force. While the bottom boundary filament remained pinned, the network was allowed to deform until *t* = 45 s, by which time networks achieve steady state (Figure S1a and Figure S1c). At *t* = 45 s, the position of each fiber in the network was recorded as the final state of the network. Center-of-mass network displacements were computed by averaging the positions of all network fibers at times *t* = 5 s and 45 s and subtracting final position from initial position. Boundary filaments were excluded from center-of-mass computations. For shear, we used the horizontal (*x*) component of displacement to estimate an elastic modulus, and in extension we used the vertical (*y*) component. For representative simulations, see Movie M1–M4 (shear force) and Movie M5–M8 (tensile force)^2^. Center-of-mass positions of these networks over time are plotted in Figure S1a and Figure S1c.

### Estimation of elastic moduli

To quantify network rigidity, we subjected each sampled network to a series of applied forces and computed the corresponding center-of-mass displacements. We generated *N* random seeds for Cytosim’s random number generator, corresponding to *N* unique initial geometries at each astral number. Specification of random seeds in the Cytosim `config.cym` file thus enabled us to apply multiple forces to the same network. For each network, we perform a sweep over *k* equally spaced force magnitudes from 0 pN to 5 pN (shear) or 0 pN to 20 pN (tensile). We then construct a force-displacement curve and perform linear regression (with intercept fixed at (0, 0)) to estimate an elastic modulus (Figure 1c). This fitting method is equivalent to estimating the slope of stress-strain data, since network stress is *F*_shear_*/s* (or *F*_tens_*/s*) and strain is Δ*x/s* (resp. Δ*y/s*). The number of force values used per network (*k*) is specified in each figure caption.

Note that the regression is performed on all (force, displacement) pairs for each network even when curvature is suggested by the data samples (see Figure S1b and Figure S1d). Also, though fractured/disconnected networks correspond to a modulus identically equal to 0, we did not attempt to detect if a network was disconnected before assigning a modulus. Instead, as fractured network components that are connected to the top boundary continue moving away from their initial positions at a near-constant speed during the force application phase (5 s *≤ t ≤* 45 s), fractured networks appear in the data as curves with remarkably high displacements. Thus, performing regression on these points produces a modulus estimate very near zero. In the small number of cases where regression yielded a negative slope (likely due to failure of the network to connect with the top boundary, thus leaving the network subject only to thermal fluctuations), a modulus of 0 was manually assigned.

### Dangling ends measurements and geometric percolation analysis

To study geometric properties of astral networks, we generated astral networks by distributing asters uniformly at random in a square domain and placing crosslinks wherever filaments intersect. Generation and analysis code^3^ is written in MATLAB R2024b (The Mathworks). All measurements of dangling ends as well as counts of productive segments by node were conducted for networks with *ρ* = 7.5 µm^−1^, ℓ = 1 µm, and *s* = 10 µm. Network parameters for percolation probability estimates are provided in the relevant figure captions.

To analyze percolation in astral networks, we used the MATLAB function `conncomp` to identify sets of mutually crosslinked filaments, henceforth referred to as connected components. In the language of graph theory, each network filament is represented by a graph vertex and an edge is drawn between vertices if those filaments were crosslinked together (i.e., if they intersected). A connected component was said to be *spanning* if it contained both a crosslink above and below the network boundaries (see Figure 5a), and an entire network was said to be spanning if it contained at least one spanning component. For *connectivity* percolation, we determined if a network had a unique connected component (see Figure 5b).

We define the critical percolation densities 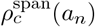 and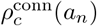 to be the network density at which the percolation probability of networks with astral number *a*_*n*_ equals 50%. To compute these values, we used the MATLAB function `fit` to compute smoothing spline fits (automatically selected smoothing parameter) to the percolation data. Using these fits, we numerically solved (using MATLAB’s `fsolve`, default tolerances) for the network density where the percolation probability reaches 50%. Example percolation data and fits are shown in Figure S6.

## Acknowledgements

We thank Shannon McFadden and Andrew Rusli for useful discussion. This work was supported by NSF DMS 2052668 and NIH T32 GM136624.

## Footnotes

1 Forked at github.com/bcberg/cytosim-bcb, with simulation configurations at github.com/bcberg/Berg_AstralMikado

2 All Movies are available at 10.5281/zenodo.15685022

3 github.com/bcberg/Berg_AstralMikado

