## Supplemental Figures for "Astral architecture can enhance mechanical strength of cytoskeletal networks by modulating percolation thresholds"

### 1 Supplemental material

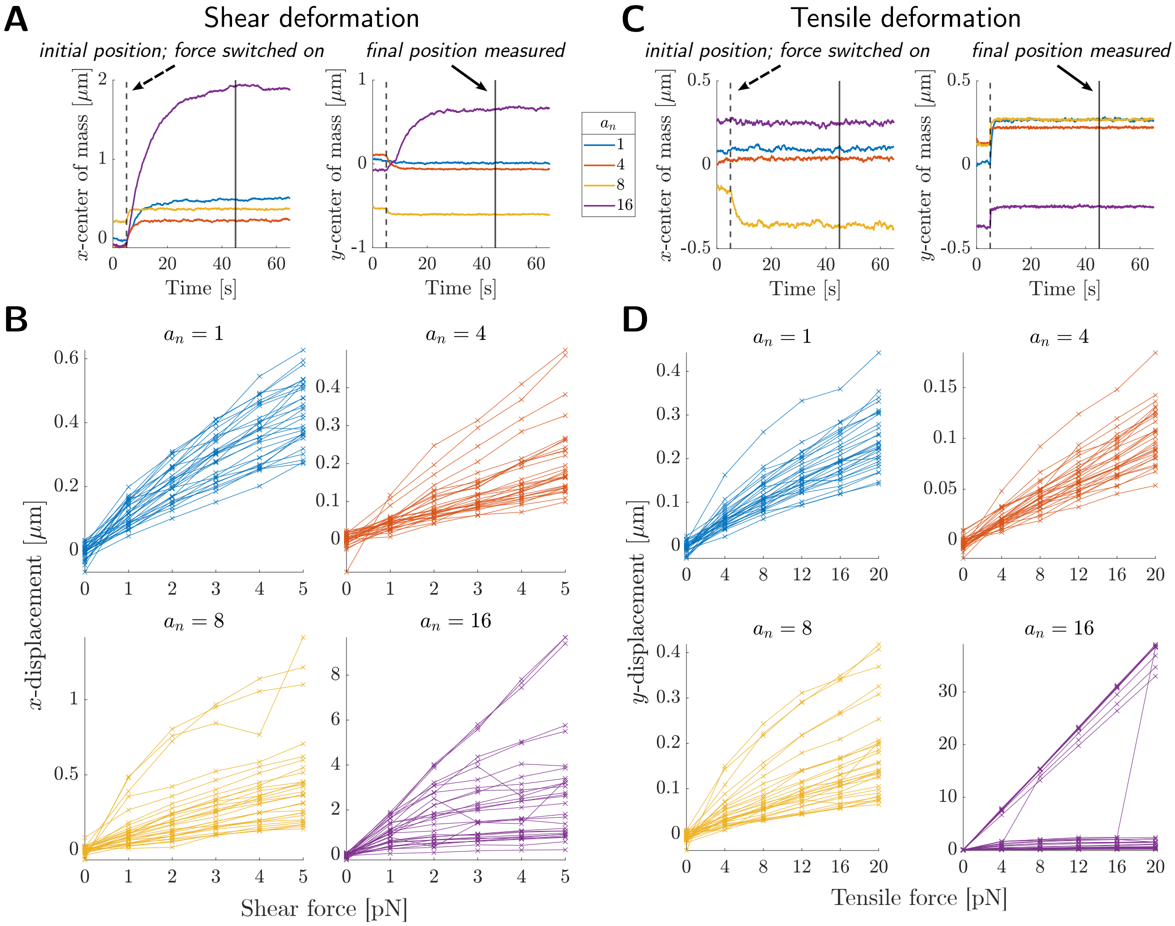


Figure S1: **Force-displacement measurements using Cytosim.** (a) Center of mass coordinates over time for individual networks experiencing a shear force of $5\,\mathrm{pN}$ (in the positive $x$-direction). Dashed vertical line indicates when the force was switched on, and solid vertical line indicates when final network positions were recorded. Legend is common to (a) and (c). Movies of these networks are provided. in M1, M2, M3, and M4. (b) Horizontal displacement data at selected astral numbers (subset of the data used to generate Figure 2b). (c) Center of mass coordinates over time for individual networks experiencing a tensile force of $20\,\mathrm{pN}$ (in the positive $y$-direction). Dashed vertical line indicates when the force was switched on, and solid vertical line indicates when final network positions were recorded. Movies of these networks are provided in M5, M6, M7, and M8. (d) Vertical displacement data at selected astral numbers (subset of the data used to generate Figure 2b). For (b) and (d), $N = 30$ networks were simulated per astral number.

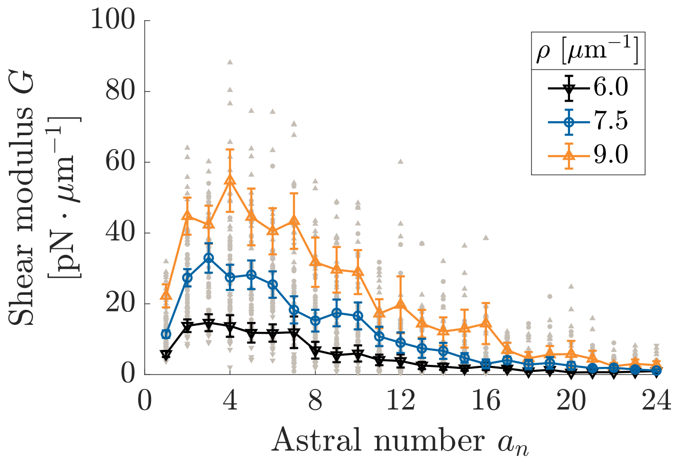


Figure S2: **Location of rigidity peak is insensitive to network density.** Shear modulus as a function of astral number for three filament densities $\rho$. Plots show mean and 95% CIs, along with individual network moduli (gray markers). The $\rho = 7.5\,\mu\text{m}^{-1}$ curve is reproduced from Figure 2b, where $N = 30$ networks were sampled per astral number with 6 force values per network. The curves at $\rho = 6\,\mu\text{m}^{-1}$ and $\rho = 9\,\mu\text{m}^{-1}$ were generated from $N = 15$ network samples per astral number and 4 force values per network.

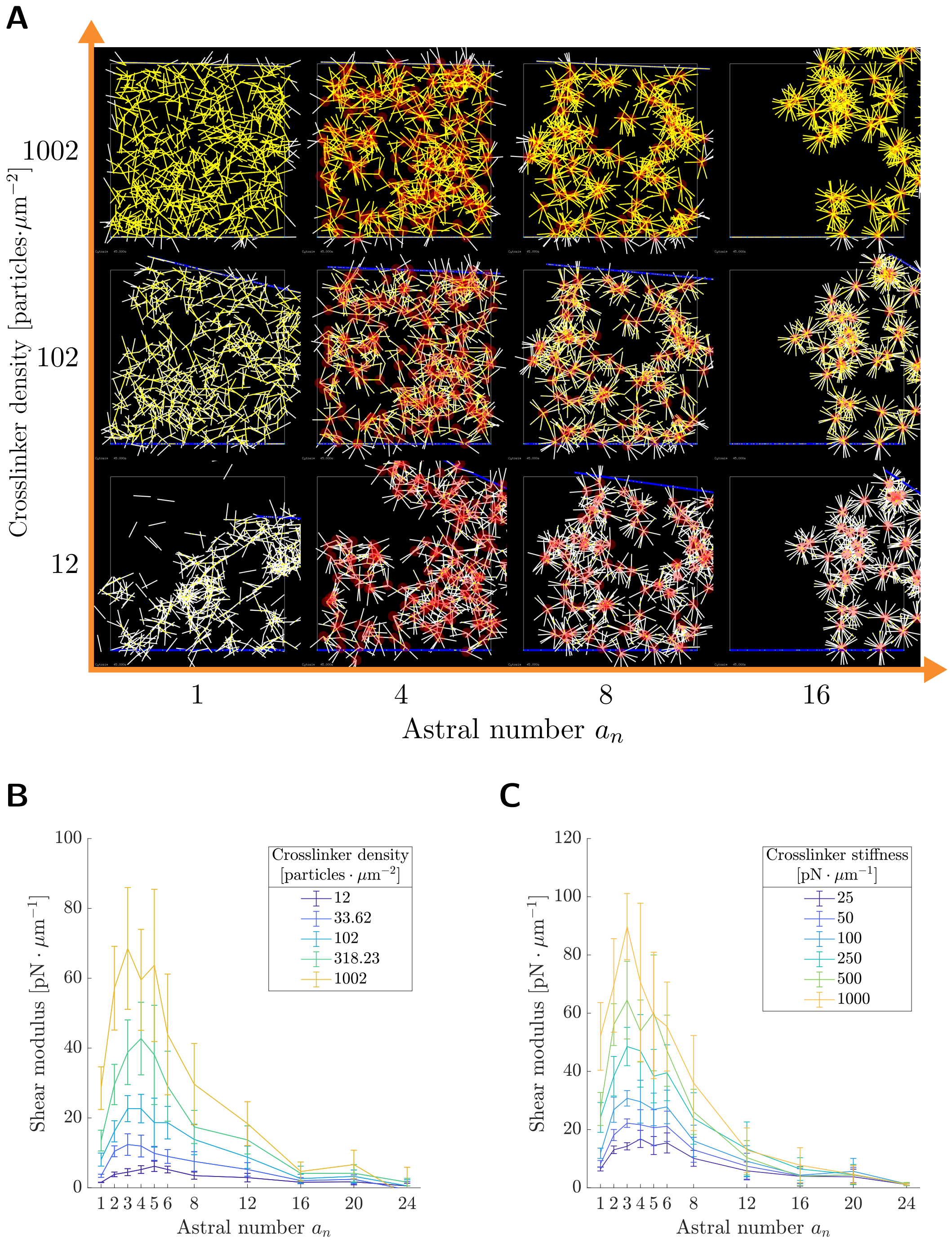


Figure S3: **Rigidity scales similarly with crosslinker density and crosslinker stiffness.** (a) Steady-state snapshots of astral network deformation at various crosslinker densities. Crosslinker particles are shown in yellow, and astral centers are marked in red. All networks have filament density $\rho = 7.5\,\mu\mathrm{m}^{-1}$ and experience a shear force of magnitude $5\,\mathrm{pN}$. (b) Shear modulus as a function of astral number for a series of crosslinker densities. (c) Shear modulus as a function of astral number for a series of crosslinker stiffnesses. Note that "stiffness" refers to the linear stretch resistance of each crosslinker, and crosslinkers do not individually exert torques. Plots show mean and 95% CIs computed from $N = 10$ networks per astral number and 4 force values per network.

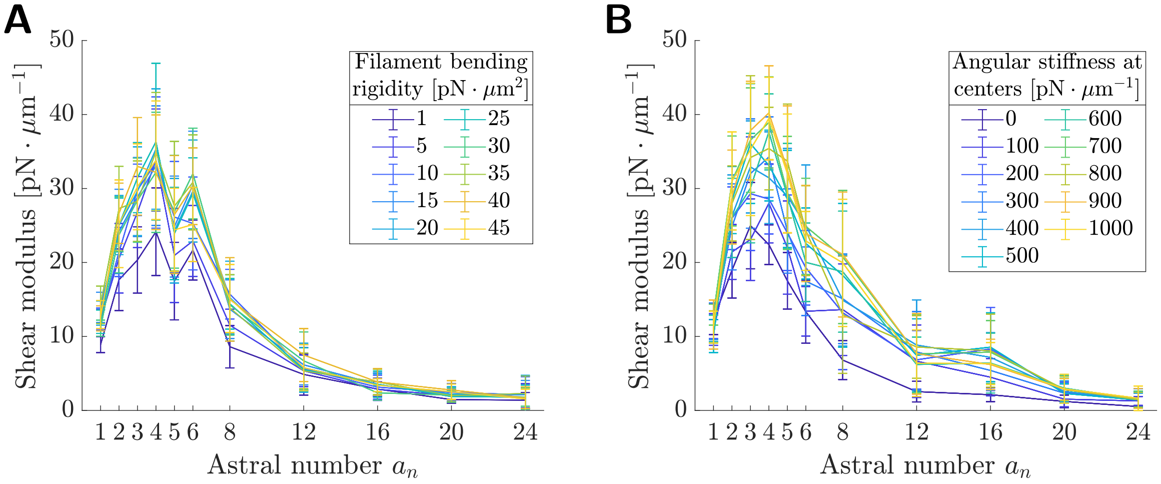


Figure S4: **Peak in rigidity is only weakly sensitive to bending of filaments and angular stiffness at astral centers.** (a) Shear modulus as a function of astral number for a series of filament bending rigidities. A default filament bending rigidity of $20\,\mathrm{pN\,\mu m^2}$ was used outside of this panel. (b) Shear modulus as a function of astral number for a series of angular stiffnesses at astral centers. A default angular stiffness of $500\,\mathrm{pN\,\mu m^{-1}}$ was used outside of this panel. Plots show mean and 95% CIs computed from $N = 10$ networks per astral number and 4 force values per network.

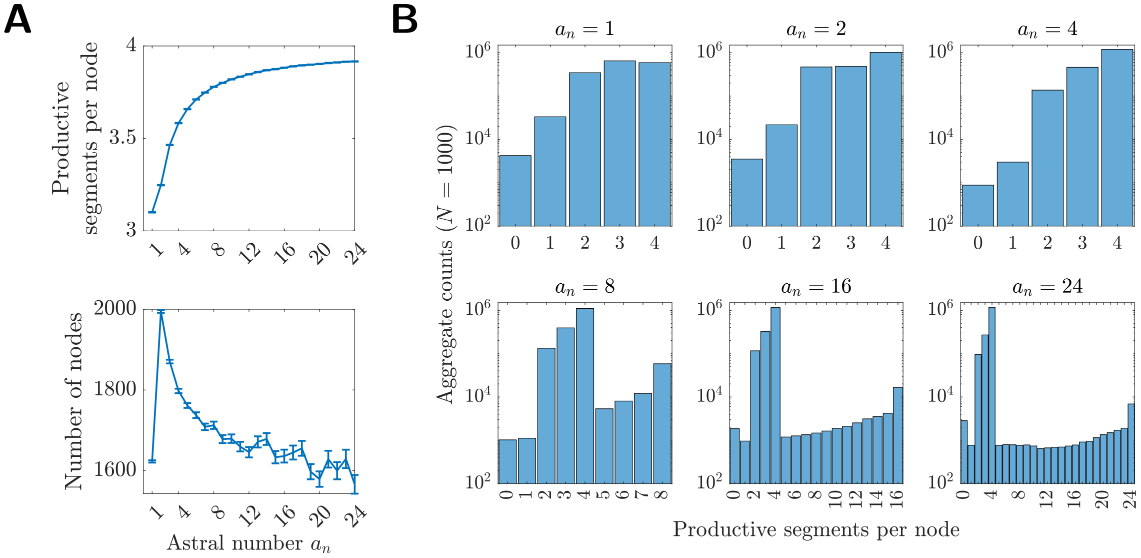


Figure S5: **Number of segments per node, including both astral centers and inter-aster crosslinks.** (a) Upper: Number of productive (i.e. non-dangling) segments per node as a function of astral number. Lower: Number of nodes (crosslinks) in astral networks as a function of astral number. Data shown are mean and 95% CIs estimated from 1000 networks per astral number. (b) Nodes from 1000 astral networks sorted by the number of productive segments connected to each node.

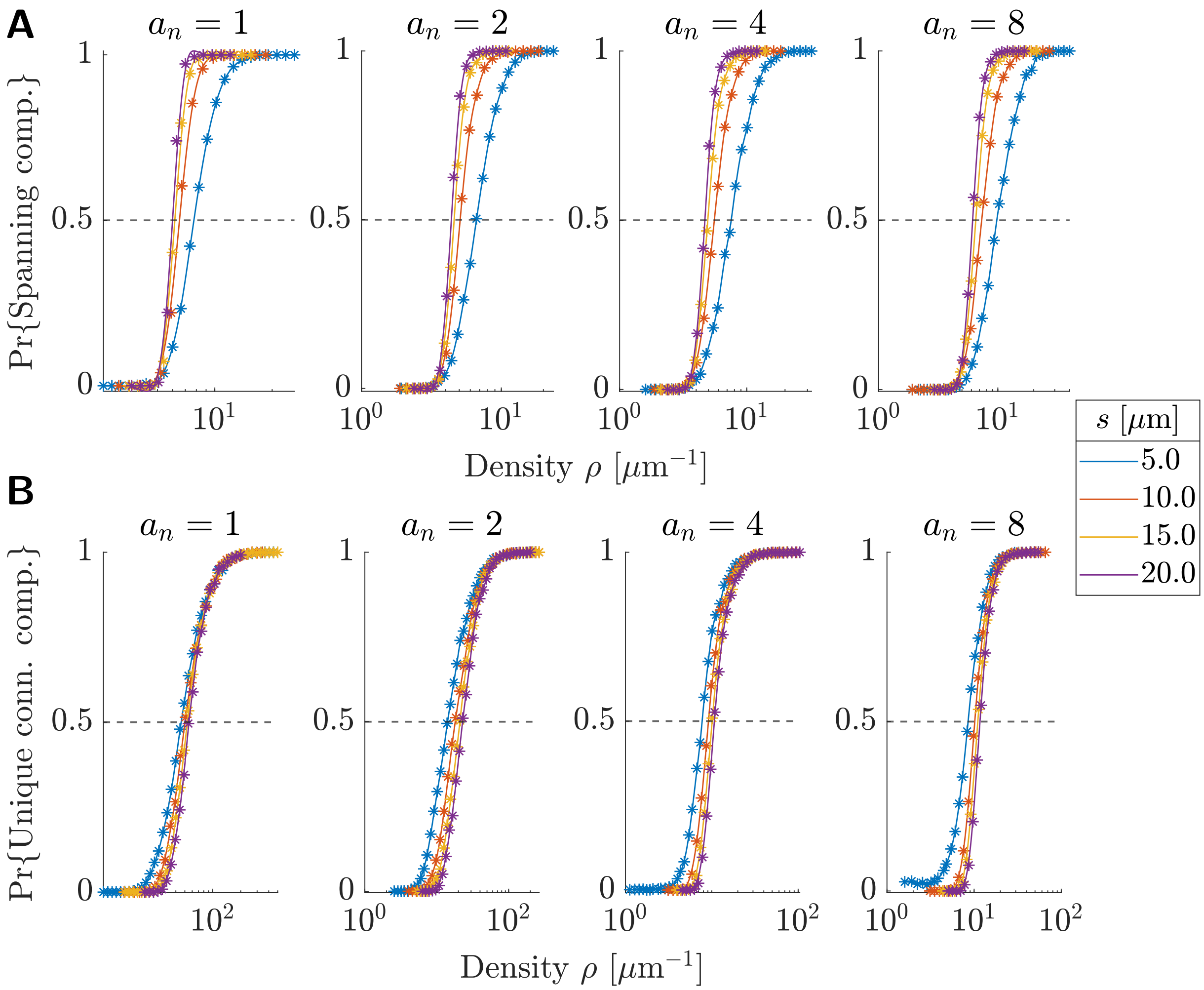


Figure S6: **Data and fits used to estimate critical percolation densities.** (a) Probability that a network contains a spanning component as a function of network density $\rho$. (b) Probability that a network contains a unique connected component as a function of network density $\rho$. Markers show percolation probabilities estimated from $N = 2000$ networks; curves show smoothing spline fits used to estimate critical percolation densities (intersection with $p = 0.5$). Data shown for selected astral numbers $a_n$ and a family of system sizes $s$. All network filaments have length $\ell = 1\,\mu$m. Only data for $s = 10\,\mu$m were used to generate Figure 5d.

#### 2 Supplemental Movies

All Movies are available at 10.5281/zenodo.15685022.

Figure M1: Simulation of a filament network with astral number $a_n = 1$ (i.e., a classical "Mikado" network) experiencing a shear force. For times $0 \leq t < 5\,\mathrm{sec}$, the network crosslinks into its initial position while no external force is applied. At time $t = 5\,\mathrm{sec}$, a constant shear force of magnitude $5\,\mathrm{pN}$ is switched on and maintained for the remainder of the simulation.

Figure M2: Simulation of an astral filament network with astral number $a_n = 4$ experiencing a shear force. For times $0 \leq t < 5\,\mathrm{sec}$, the network crosslinks into its initial position while no external force is applied. At time $t = 5\,\mathrm{sec}$, a constant shear force of magnitude $5\,\mathrm{pN}$ is switched on and maintained for the remainder of the simulation.

Figure M3: Simulation of an astral filament network with astral number $a_n = 8$ experiencing a shear force. For times $0 \leq t < 5\,\mathrm{sec}$, the network crosslinks into its initial position while no external force is applied. At time $t = 5\,\mathrm{sec}$, a constant shear force of magnitude $5\,\mathrm{pN}$ is switched on and maintained for the remainder of the simulation.

Figure M4: Simulation of an astral filament network with astral number $a_n = 16$ experiencing a shear force. For times $0 \leq t < 5\,\mathrm{sec}$, the network crosslinks into its initial position while no external force is applied. At time $t = 5\,\mathrm{sec}$, a constant shear force of magnitude $5\,\mathrm{pN}$ is switched on and maintained for the remainder of the simulation.

Figure M5: Simulation of a filament network with astral number $a_n = 1$ (i.e., a classical "Mikado" network) experiencing a tensile force. For times $0 \leq t < 5\,\mathrm{sec}$, the network crosslinks into its initial position while no external force is applied. At time $t = 5\,\mathrm{sec}$, a constant tensile force of magnitude $20\,\mathrm{pN}$ is switched on and maintained for the remainder of the simulation.

Figure M6: Simulation of an astral filament network with astral number $a_n = 4$ experiencing a tensile force. For times $0 \leq t < 5\,\mathrm{sec}$, the network crosslinks into its initial position while no external force is applied. At time $t = 5\,\mathrm{sec}$, a constant tensile force of magnitude $20\,\mathrm{pN}$ is switched on and maintained for the remainder of the simulation.

Figure M7: Simulation of an astral filament network with astral number $a_n = 8$ experiencing a tensile force. For times $0 \leq t < 5\,\mathrm{sec}$, the network crosslinks into its initial position while no external force is applied. At time $t = 5\,\mathrm{sec}$, a constant tensile force of magnitude $20\,\mathrm{pN}$ is switched on and maintained for the remainder of the simulation.

Figure M8: Simulation of an astral filament network with astral number $a_n = 16$ experiencing a tensile force. For times $0 \leq t < 5\,\mathrm{sec}$, the network crosslinks into its initial position while no external force is applied. At time $t = 5\,\mathrm{sec}$, a constant tensile force of magnitude $20\,\mathrm{pN}$ is switched on and maintained for the remainder of the simulation.
